# Lack of insurance is associated with lower probability of diagnostic imaging use among US trauma patients: An instrumental variable analysis and simulation

**DOI:** 10.1101/215889

**Authors:** Audrey Renson, Finn D. Schubert, Marc A. Bjurlin

## Abstract

**Background:** Uninsured trauma patients have higher mortality than their insured counterparts. One possible reason is disparities in utilization of appropriate diagnostic imaging, including computed tomography (CT), X-ray, ultrasound (US), and magnetic resonance imaging (MRI). We examined the association between lack of insurance and use of diagnostic imaging.

**Methods:** Data come from the National Trauma Databank 2010-2015. Patients were determined uninsured if payment mode was self-pay or missing. The primary outcome was any diagnostic imaging procedure, and secondary outcomes included CT, X-ray, US, or MRI. Risk ratios (RRs) were adjusted for demographics, comorbidities, injury characteristics, facility characteristics. We also used the 2010 Patient Protection and Affordable Care Act as an instrumental variable (IV), with linear terms for year to account for annual trends in imaging use. Monte carlo simulations to test effect of hypothetical violations to IV assumptions of relevance, no direct effect, and no confounding.

**Results:** Of 4,373,554 patients, 953,281 (21.8%) were uninsured. After adjusting, uninsured patients had lower chance of any imaging (RR 0.98, 95% CI 0.98 to 0.98), x-ray (RR 0.99, 95% CI 0.99 to 1.00), and MRI (RR 0.82, 95% CI 0.81 to 0.83), and higher chance of ultrasound (RR 1.01, 95% CI 1.01 to 1.02). In IV analysis, uninsured status was associated with reduction in any imaging (RR 0.60, 95% CI 0.52 to 0.70), tomography (RR 0.52, 95% CI 0.44 to 0.62) ultrasound (RR 0.46, 95% CI 0.32 to 0.65), and MRI (RR 0.19, 95% CI 0.10 to 0.37) and increased likelihood of x-ray use (RR 1.74, 95% CI 1.31 to 2.32). Simulations indicated that a direct effect RD of −0.02 would be necessary to produce observed results under the null hypothesis.

**Discussion:** Our study suggests an association between insurance status and use of imaging that is unlikely to be driven by confounding or violations of IV assumptions. Mechanisms for this remain unclear, but could include unconscious provider bias or institutional financial constraints. Further research is warranted to elucidate mechanisms and assess whether differences in diagnostic imaging use mediate the association between insurance and mortality.

## INTRODUCTION

Health inequalities based on health insurance status have long been observed. In particular, US adults without insurance have a higher rate of mortality,^1^ with lack of insurance estimated to cost tens of thousands of lives per year in the US.^2^ Insured patients generally have better self-rated health than non-insured patients,^3,4^ likely related to greater access to primary care and increased use of preventative services.^5^

Given that unintentional injury is the 3^rd^ leading cause of death in the US,^6^ health insurance-related outcome disparities among trauma patients are of significant concern. In trauma patients specifically, with rare exceptions,^7^ not having health insurance has been associated with negative outcomes, such as failure-to-rescue^8,9^ and hospital mortality,^10-16^ as well as two-year mortality.^17^ Hospital-based studies of trauma patients have also found disparities in post-hospitalization care associated with insurance status.^18,19^

One possible mechanism for health insurance-related disparities in trauma patients is differences in care, including receipt of fewer diagnostic tests. The Emergency Medical Treatment and Active Labor Act (1986) explicitly prohibits transfer or refusal of necessary treatment in a medically unstable patient, and is intended to prevent discrimination in care decisions based on insurance status.^20^ Despite this, in a systematic review, uninsured critically ill patients had 8.5% fewer procedures and were more likely to experience hospital discharge delays and to have life support withdrawn, compared to their insured counterparts.^21^ Being uninsured has also been associated with decreased odds of receiving a central venous catheter, acute hemodialysis, and tracheostomy.^12^ Among patients with pelvic fractures, uninsured patients have been found to receive fewer diagnostic procedures, with a larger disparity for more resource-intensive tests.^22^

Prompt diagnosis and triage via appropriate diagnostic imaging, including computed tomography (CT),^23-25^ X-ray radiography,^26-28^ ultrasound (US),^29-31^ and magnetic resonance imaging (MRI),^24,25,32^ plays a key role in initial management of traumatic injuries and can decrease mortality. However, to our knowledge, no published studies have examined health insurance-related disparities in the use of diagnostic imaging procedures among trauma patients in general. Our study uses data from a national sample of trauma patients to estimate the impact of insurance status on use of diagnostic imaging procedures.

## METHODS

### Study Population and Definitions

#### Data Source

This study used data from years 2010 through 2015 of the National Trauma Data Bank (NTDB), the largest collection of trauma registry data available in the United States. The data are submitted voluntarily by an increasing number of trauma centers in the US—682 hospitals submitted data to the NTDB in 2010,^33^ and 746 did so in 2015,^34^ comprising Level I, II, III, and IV Trauma Centers, as well as non-designated facilities, including pediatric-only centers, in all regions of the US. The NTDB includes data regarding the characteristics of the admitting facility, patient demographics, mechanism of injury and injury severity, care provided during the hospital stay, and discharge disposition.

#### Exposure and Outcome Variables

The exposure of interest was patient’s insurance status. Because the NTDB only collects payment information and not insurance status directly, we defined patients as uninsured if their payment method was marked as self-pay or was missing. The primary outcome was the use of any diagnostic imaging procedure during the hospital admission following trauma. We separately considered the use of tomography, x-ray, ultrasound, and magnetic resonance imaging (MRI) as secondary outcomes.

### Statistical Methods

#### Confounding Variables

To adjust for confounding variables, we calculated a propensity score using logistic regression consisting of age, gender, race/ethnicity, injury severity score (ISS), any and all comorbidities entered individually, along with the facility-level variables of hospital type (university, community, non-teaching), ACS trauma level, bed size, number of neuro surgeons, number of trauma surgeons, and U.S. geographical region (northeast, south, midwest, west). Predicted values from the propensity model were stratified into deciles and entered into adjusted models as nominal covariates.

#### Crude and Adjusted Models

To estimate the association of being uninsured with probability of any imaging, tomography, x-ray, ultrasound, and MRI, we calculated both risk differences and risk ratios. For each measure, we calculated crude as well as adjusted measures. Risk differences were calculated with linear regression (also known as a linear probability model), and risk ratios were calculated using Poisson regression with robust sandwich standard error estimates.^35^

#### Instrumental Variable Analysis

The impact of patient insurance status on imaging use is likely to be confounded by patient- and area-level socioeconomic status (SES), variables which are not measured in the NTDB. To address unmeasured confounding, we used instrumental variables (IV) analysis, with the 2010 US Patient Protection and Affordable Care Act (ACA) coverage provisions as an instrument. In the context of IV analysis, an instrument is a variable *Z* meeting three criteria: (i) relevance, i.e. that *Z* has a causal effect on the outcome, *Y;* (ii) the exclusion restriction, i.e. that *Z* has no effect on *Y* through any pathway except the putative exposure, *X*; and (iii) no confounding, i.e. *Z* and *Y* do not share any common causes. ^36^

With certain caveats, the ACA represents an instrument, in that as of January 1, 2014, expansion of the Medicaid program and federal subsidies for private insurance under the ACA directly caused patients to obtain insurance, and patients shortly before and after this date would be similar in almost all ways except for having insurance. A potential violation of (ii) is that changes in Medicare and Medicaid reimbursement rules introduced by the ACA served to de-incentivize uncritical use of diagnostic imaging services,^37^ as did the US Deficit Reduction Act of 2005,^38^ which would bias in the opposite direction of our hypothesis. A countervailing trend and potential violation of (iii) is the rapid advancement occurring in radiology technology, such that imaging use has been increasing at the same time as ACA changes to insurance ^39-41^. This would bias in the direction of our hypothesis, but is a steady rather than abrupt effect that can be addressed by de-trending of the data prior to treating the January 1, 2014 cutoff as an instrument.

For each outcome, we used two-stage least squares to calculate IV risk differences (RDs), and two-stage logistic regression method modified by using a log (instead of logit) link in second stage, to calculate IV risk ratios (RRs). Both two-stage least squares and two-stage logistic regression have been shown to be unbiased for binary instrument, exposure, and response variables;^42^ a log link in the second stage should not in principle introduce any bias, and is necessary to estimate the relative risk in the case of a common outcome such as imaging. Both first and second stage models in both approaches included a linear term for year to address any constant yearly increase in imaging use.

#### Sensitivity Analysis

Assumptions (ii) and (iii) of the IV method are unverifiable and contain some potential for violation in our study (especially (ii) – the addition of the year term should remove the violation to (iii)). Further, our instrument was relatively weak (0.023 on the RD scale), which can induce large variance in effect measures even when no conditions are violated, and will amplify bias in the case that (ii) and/or (iii) are violated.^36^ We therefore conducted sensitivity analyses to determine whether the weakness of our instrument alone could produce the observed results, or if not, the size of direct effect (ii) and/or unmeasured confounding (iii) that would be necessary to produce them, given the null hypothesis of no effect of insurance on imaging rates.

To do this, we simulated datasets under three scenarios. In the first scenario, IV conditions were met by design, and the strength of the instrument was varied from 0 to 0.3 on the RD scale. In the second scenario, the strength of the instrument was set at that observed in our data (RD=0.023), and condition (ii) was violated with direct effects ranging from 0 to 0.3 on the RD scale, with no violation to condition (iii). In the third scenario, the strength of the instrument was again set at RD=0.023, condition (ii) was met by design, and condition (iii) was violated with bias due to confounding ranging from 0 to 0.3 in terms of difference in RD. For each dataset, we create 10,000 observations and with 50 replications with different random seeds. For each measure of interest, we calculated the measure on each of the 50 replicated datasets and took the mean value.

All statistical analysis were conducted using R software version 3.3.2,^43^ with instrumental variable analysis carried out using the packages ‘AER’^44^ and ‘ivpack’,^45^ and simulation data generated using the package ‘simstudy’.^46^

## RESULTS

The 2010-2015 NTDB sample consisted 4,936,880 patient records, of whom 563,326 were excluded because their payment mode was coded as no fault automobile, workers compensation, not billed, or other. This left 4,373,554 patient records, of whom 953,281 (21.8%) were classified as uninsured and 3,420,273 (78.2%) as insured. Half the sample (50.0%) had at least one imaging procedure, the most frequent of which was tomography (39.7%), followed by x-ray (20.1%), ultrasound (15.0%), and MRI (4.5%). (Table 1) Figure 1 illustrates trends in imaging and rates of uninsured over time. In general, between 2010 and 2015, imaging use was increasing, with the strongest trend demonstrated in tomography, and the smallest increase in MRI use. Conversely, the rate of uninsured was decreasing, with a marked change occurring between 2013 and 2014.

**Figure 1.**
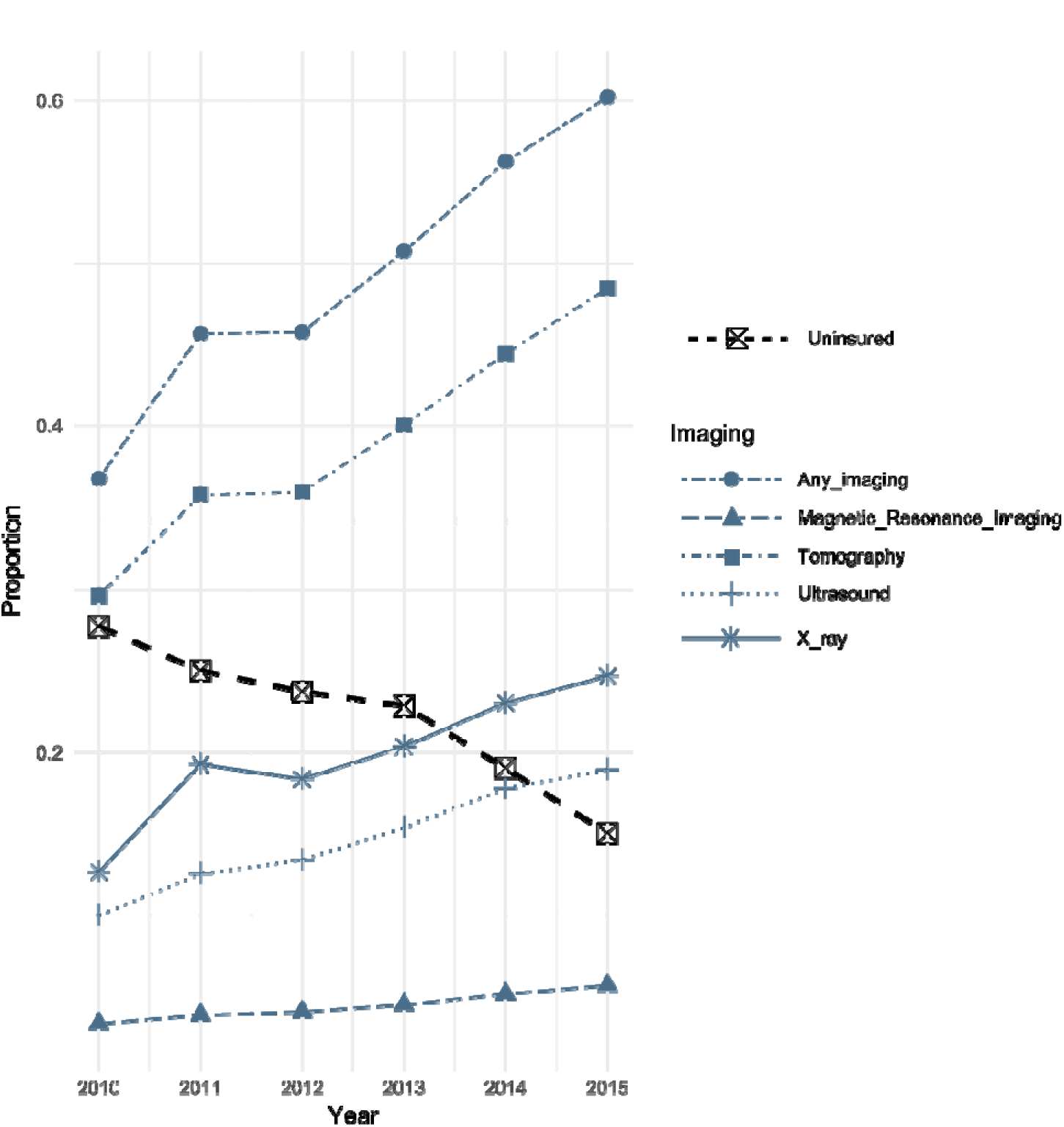
Yearly trends in imaging and uninsured status, in all patients reported to the US National Trauma Data Bank, 2010-2015, excluding patients whose payment mode was coded as no-fault automobile, workers compensation, not billed, or other.

**Table 1.** Descriptive statistics.

|  | Overall | Uninsured | Insured | P-value |
| --- | --- | --- | --- | --- |
| Total | 4,373,554 | 953,281 | 3,420,273 |  |
| Age (years) -- Median [IQR] | 44.0 [23.0 to 66.0] | 34.0 [24.0 to 49.0] | 50.0 [22.0 to 71.0] | <0.001 |
| ISS -- Median [IQR] | 9.0 [ 4.0 to 13.0] | 6.0 [ 4.0 to 13.0] | 9.0 [ 4.0 to 13.0] | <0.001 |
| Gender=Male -- n (%) | 2,670,207 (61.1%) | 709,912 (74.5%) | 1,960,295 (57.3%) | <0.001 |
| Race/ethnicity -- n (%) |  |  |  | <0.001 |
| American Indian | 38,413 (0.9%) | 7,440 (0.8%) | 30,973 (0.9%) |  |
| Asian or Pacific Islander | 82,727 (2.0%) | 17,786 (2.0%) | 64,941 (2.0%) |  |
| Non-Hispanic Black | 615,549 (14.7%) | 206,337 (23.2%) | 409,212 (12.5%) |  |
| Non-Hispanic White | 3,062,680 (73.4%) | 540,517 (60.7%) | 2,522,163 (76.8%) |  |
| Other Non-Hispanic (including multi-racial) | 375,785 (9.0%) | 119,038 (13.4%) | 256,747 (7.8%) |  |
| Hospital type -- n (%) |  |  |  | <0.001 |
| Community | 1,629,071 (37.2%) | 311,960 (32.7%) | 1,317,111 (38.5%) |  |
| Non-Teaching | 705,185 (16.1%) | 145,433 (15.3%) | 559,752 (16.4%) |  |
| University | 2,039,298 (46.6%) | 495,888 (52.0%) | 1,543,410 (45.1%) |  |
| ACS Trauma Level -- n (%) |  |  |  | <0.001 |
| I | 1,415,312 (32.4%) | 379,746 (39.8%) | 1,035,566 (30.3%) |  |
| II | 884,883 (20.2%) | 174,015 (18.3%) | 710,868 (20.8%) |  |
| III | 158,365 (3.6%) | 26,733 (2.8%) | 131,632 (3.8%) |  |
| IV | 6,386 (0.1%) | 1,852 (0.2%) | 4,534 (0.1%) |  |
| None reported | 1,908,608 (43.6%) | 370,935 (38.9%) | 1,537,673 (45.0%) |  |
| Bed size -- n (%) |  |  |  | <0.001 |
| <=200 | 353,311 (8.1%) | 69,863 (7.3%) | 283,448 (8.3%) |  |
| 201 - 400 | 1,298,117 (29.7%) | 249,677 (26.2%) | 1,048,440 (30.7%) |  |
| 401 - 600 | 1,230,527 (28.1%) | 271,828 (28.5%) | 958,699 (28.0%) |  |
| > 600 | 1,491,599 (34.1%) | 361,913 (38.0%) | 1,129,686 (33.0%) |  |
| Region of US -- n (%) |  |  |  | <0.001 |
| Midwest | 1,153,510 (26.6%) | 229,682 (24.3%) | 923,828 (27.2%) |  |
| Northeast | 697,808 (16.1%) | 104,420 (11.0%) | 593,388 (17.5%) |  |
| South | 1,589,157 (36.6%) | 416,663 (44.1%) | 1,172,494 (34.5%) |  |
| West | 903,383 (20.8%) | 194,547 (20.6%) | 708,836 (20.9%) |  |
| Neuro Surgeons -- n (%) |  |  |  | <0.001 |
| 0 | 212,296 (4.9%) | 43,875 (4.6%) | 168,421 (4.9%) |  |
| 1-3 | 2,210,437 (50.8%) | 454,391 (47.8%) | 1,756,046 (51.6%) |  |
| 4-5 | 442,648 (10.2%) | 117,415 (12.4%) | 325,233 (9.6%) |  |
| 6-7 | 1,289,868 (29.6%) | 285,804 (30.1%) | 1,004,064 (29.5%) |  |
| 8-9 | 102,253 (2.3%) | 23,623 (2.5%) | 78,630 (2.3%) |  |
| 10+ | 97,508 (2.2%) | 25,299 (2.7%) | 72,209 (2.1%) |  |
| Trauma surgeons -- n (%) |  |  |  | <0.001 |
| 0 | 33,151 (0.8%) | 6,943 (0.7%) | 26,208 (0.8%) |  |
| 1-3 | 281,943 (6.4%) | 55,878 (5.9%) | 226,065 (6.6%) |  |
| 4-5 | 1,906,698 (43.6%) | 396,138 (41.6%) | 1,510,560 (44.2%) |  |
| 6-7 | 1,080,163 (24.7%) | 245,828 (25.8%) | 834,335 (24.4%) |  |
| 8+ | 1,071,599 (24.5%) | 248,494 (26.1%) | 823,105 (24.1%) |  |

IQR=interquartile range; ISS=injury severity score; ACS=American College of Surgeons; US=United States

Table 2 displays crude, adjusted, and IV estimates of RRs and RDs for any imaging, tomography, x-ray, ultrasound, and MRI. When estimating the crude association of being uninsured with receiving imaging, uninsured patients had a lower probability of any imaging (RR 0.93, 95% CI 0.92 to 0.93), tomography (RR 0.94, 95% CI 0.94 to 0.94), and x-ray (RR 0.93, 95% CI 0.93 to 0.94). This negative association was even more pronounced in MRI (RR 0.73, 95% CI 0.73 to 0.74), whereas a significant but negligible positive association was present for ultrasound (RR 1.01, 95% CI 1.01 to 1.02). After adjusting for age, gender, race/ethnicity, ISS, comorbidities, and facility characteristics, associations were attenuated but still present with any imaging (RR 0.98, 95% CI 0.98 to 0.98), x-ray (RR 0.99, 95% CI 0.99 to 1.00), and MRI (RR 0.82, 95% CI 0.81 to 0.83), but no longer present for tomography (RR 1.00, 95% CI 1.00 to 1.00). The slight positive association with ultrasound use remained unchanged after adjustment (RR 1.01, 95% CI 1.01 to 1.02).

**Table 2.** Association of being uninsured on probability of any imaging procedure.

|  | Crude |  |  | Adjusted<sup>a</sup> |  |  | IV<sup>b</sup> |  |  |
| --- | --- | --- | --- | --- | --- | --- | --- | --- | --- |
|  | Estimate | 95% CI | P-value | Estimate | 95% CI | P-value | Estimate | 95% CI | P-value |
| **Risk Differences** |  |  |  |  |  |  |  |  |  |
| Any imaging | -0.04 | -0.04 to -0.04 | <0.0001 | -0.01 | -0.01 to -0.01 | <0.0001 | -0.46 | -0.54 to -0.38 | <0.0001 |
| Tomography | -0.02 | -0.03 to -0.02 | <0.0001 | -0.00 | -0.00 to 0.00 | 0.231 | -0.62 | -0.71 to -0.54 | <0.0001 |
| X ray | -0.01 | -0.02 to -0.01 | <0.0001 | -0.00 | -0.00 to -0.00 | 0.002 | 0.08 | 0.02 to 0.14 | 0.006 |
| Ultrasound | 0.00 | 0.00 to 0.00 | <0.0001 | 0.00 | 0.00 to 0.00 | <0.0001 | -0.25 | -0.31 to -0.20 | <0.0001 |
| MRI | -0.01 | -0.01 to -0.01 | <0.0001 | -0.01 | -0.01 to -0.01 | <0.0001 | -0.14 | -0.17 to -0.11 | <0.0001 |
| **Risk Ratios** |  |  |  |  |  |  |  |  |  |
| Any imaging | 0.93 | 0.92 to 0.93 | <0.0001 | 0.98 | 0.98 to 0.98 | <0.0001 | 0.60 | 0.52 to 0.70 | <0.001 |
| Tomography | 0.94 | 0.94 to 0.94 | <0.0001 | 1.00 | 1.00 to 1.00 | 0.434 | 0.52 | 0.44 to 0.62 | <0.001 |
| X ray | 0.93 | 0.93 to 0.94 | <0.0001 | 0.99 | 0.99 to 1.00 | 0.007 | 1.74 | 1.31 to 2.32 | <0.001 |
| Ultrasound | 1.01 | 1.01 to 1.02 | <0.0001 | 1.01 | 1.01 to 1.02 | <0.0001 | 0.46 | 0.32 to 0.65 | <0.001 |
| MRI | 0.73 | 0.73 to 0.74 | <0.0001 | 0.82 | 0.81 to 0.83 | <0.0001 | 0.19 | 0.10 to 0.37 | <0.001 |

<sup>a</sup>Adjusted for age, gender, race/ethnicity, injury severity score (ISS), any and all comorbidities reported (each treated separately), facility type (university, community, non-teaching), ACS trauma level, facility bed size, U.S. geographic region, number of neuro surgeons, and number of trauma surgeons.
<sup>b</sup>IV risk differences calculated using two-stage least squares, risk ratios calculated using two-stage logistic regression method modified by using a log (instead of logit) link in second stage
CI=confidence interval; IV=instrumental variable; MRI=magnetic resonance imaging

When examining IV estimates, the association between uninsured status and imaging was markedly different than crude and adjusted estimates. On the whole, uninsured status was associated with a far lower probability of any imaging (RR 0.60, 95% CI 0.52 to 0.70), along with lower rates of tomography (RR 0.52, 95% CI 0.44 to 0.62) and ultrasound (RR 0.46, 95% CI 0.32 to 0.65), with the most pronounced negative association again occurring in MRI (RR 0.19, 95% CI 0.10 to 0.37). Conversely, uninsured status was associated with a greater likelihood of x-ray use (RR 1.74, 95% CI 1.31 to 2.32).

### Simulations

Figure 2a shows the results of simulating a range of instrument strengths (between 0.5 and 5 per 100 on the risk difference (RD) scale), under the assumption that no IV conditions were violated, with effect sizes compared to crude, adjusted, and true RDs assumed to be null. Given a sample size of 4,373,554 and the same degree of measured confounding observed in our study, the IV estimate greatly exaggerates the RD when the instrument is very weak (0.5 per 100), but offers an improvement over the adjusted RD when the instrument strength reaches 1 per 100. With an instrument strength of 2.33 per 100 as observed in our study, the IV risk difference is nearly equivalent to true RD, indicating that variance due to a weak instrument alone is highly unlikely to be responsible for the associations we observed.

In Figure 2b, condition (ii) was violated by introducing a direct effect of the IV on the response, varying the direct effect RD from 0 to −1.8 per 100, while holding constant the instrument strength at 2.33 per 100, and assuming no violation of condition (iii). We observe that under these circumstances, the IV RD departs rapidly from the true effect, and assuming the null hypothesis, a direct effect RD of - 1.65 per 100 would be sufficient to generate our observed IV RD of 0.60 for any imaging.

In Figure 2c, we violated condition (iii) by introducing an unmeasured confounder of the association between the IV and the response, varying the amount of bias from 0 to 1.5 per 100, measuring the departure from the true RD of the effect of the IV on the response. In this simulation, we again held the instrument strength constant at 2.33 per 100 and assumed no violation of condition (ii). Here, we again observe that the IV RD departs rapidly from the true effect, and assuming the null hypothesis, a relatively small amount of bias (1.60 per 100) would be sufficient to generate our observed IV RD of 0.60 for any imaging.

**Figure 2.**
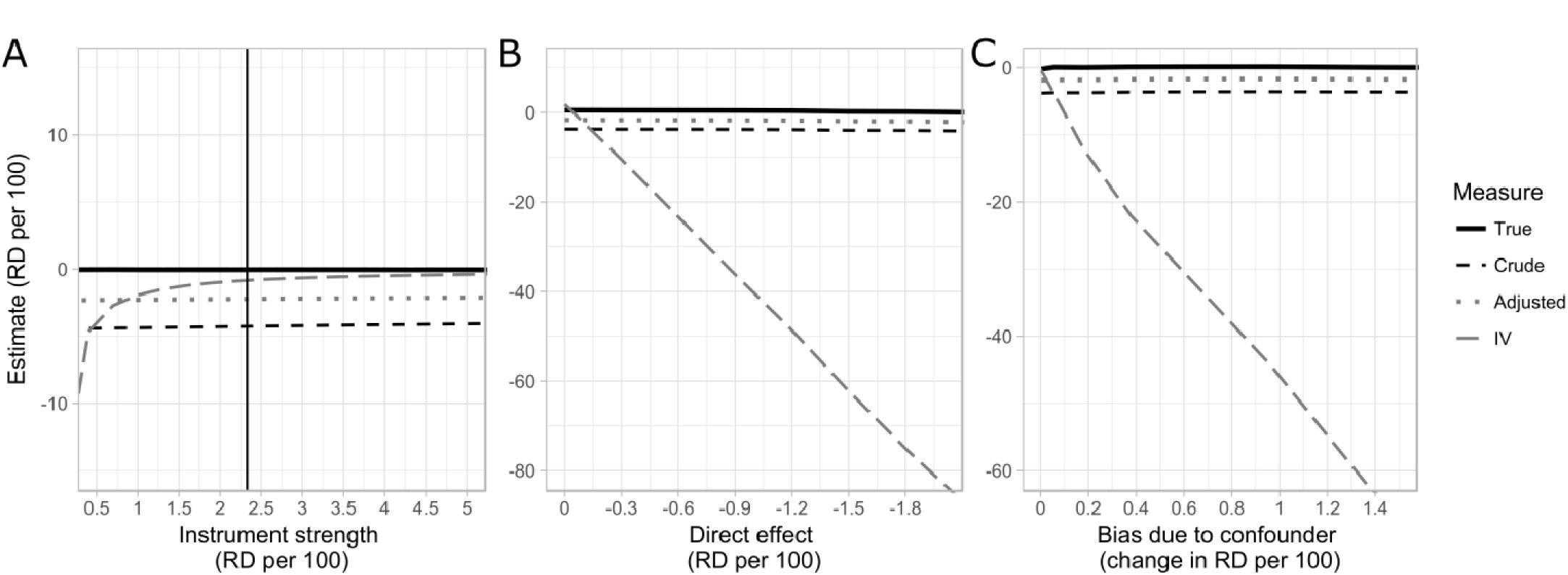
Results of simulations examining the possible impact of violation of instrumental variable assumptions, using 10 repeats f 50,000 Monte Carlo simulated observations. In each simulation, observed confounding is generated equivalently to that found in the analysis of NTDB data for our study, and an unobserved confounder is generated of similar strength. In (A), weak instrument bias is examined by varying the instrument strength and maintaining conditions (ii) and (iii), of no direct effect and no confounding of the effect of instrument on outcome, respectively. In (B), the instrument strength is held constant at 2.33 per 100 and a direct effect of varying strength is introduced. In (C), the instrument strength is held constant at 2.33 per 100 and a confounder of the effect of instrument on outcome of varying strength is introduced.

## DISCUSSION

In a national sample of over 4 million trauma patients, we found in crude analysis that uninsured patients had a lower probability of having had any imaging, with the strongest association seen between being uninsured and not having had an MRI. Lower probability of imaging among uninsured patients remained consistent even after adjusting for age, gender, race/ethnicity, ISS, comorbidities, and facility characteristics, although associations were attenuated for any imaging, X-ray, and MRI. In contrast, in both crude and adjusted analysis, uninsured patients had a slightly greater likelihood of receiving ultrasound imaging.

Using the pre- vs. post-ACA period as an instrumental variable, we found similarly that uninsured status was associated with a substantially lower probability of any imaging, tomography, ultrasound, and MRI, with MRI associated with the largest reduction in probability. Uninsured status, however, was associated with a greater likelihood of x-ray use in IV analysis.

Our study also explored the potential for bias in the IV analysis. This analysis may be affected by bias due to a direct effect of the ACA on reduction in imaging use via changes in Medicare and Medicaid reimbursement policies, a bias which would be amplified by the relative weakness of the instrument. We found in simulations that a moderate direct effect RD of approximately −1.00 per 100 would be sufficient to generate the observed IV RD of −0.46 per 100, if no true association were present. However, the plausible direct effect of the ACA would bias in the opposite direction of the observed results, in that de-incentivized imaging under the ACA would make imaging of insured patients *less* likely, whereas our results indicate the reverse. An additional possible bias in the IV analysis is confounding of the IV-outcome relationship by time, in the form of a trend towards greater imaging use over time, which plausibly should have been a steady, approximately linear yearly increase, which was eliminated by the inclusion of a linear term for year in the IV analysis. Therefore, it is unlikely that violation of IV conditions alone generated the observed association, and our results suggest a relationship between uninsured status and reduced imaging use.

These results are consistent with previous NTDB studies that focused on insurance-related disparities in hospital care in various subsets of trauma patients. One study found that uninsured patients with pelvic fractures were more likely to receive radiographs and less likely to receive vascular ultrasonography and CT of the abdomen.^47^ Another NTDB study, looking at procedural volume among traumatic brain injury patients, found that lack of insurance was associated with lower procedural volume,^48^ and this result is consistent with an NTDB analysis that found that spinal cord-injured patients with insurance had a higher probability of surgery than those without insurance.^49^ Given that many studies have revealed higher mortality rates in uninsured trauma patients,^10-15,17^ it appears plausible that one mechanism for these increased mortality rates may be disparities in diagnostic and procedural care provided during the hospital stay. Further studies should explore the relationship between disparities in hospital care and disparities in outcomes to evaluate the extent to which disparities in hospital care explain the differences seen in outcomes.

Our study expands on previous studies on insurance-related disparities in hospital care by looking at a broad sample of trauma patients, rather than limiting analysis to a specific subset of diagnoses, and found significant differences in probability of diagnostic imaging related to insurance status using two different analytic techniques, providing further illustration of insurance-related disparities in care among trauma patients.

One potential mechanism for this association is unconscious bias among providers. Previous studies among trauma surgeons and acute care physicians have demonstrated that a majority had an Implicit Association Test (IAT) score demonstrating an unconscious preference toward white persons and persons of upper social class, however, IAT scores were not associated with responses to clinical vignettes.^50^ Despite this, there is some evidence that even when physicians deny bias, treatment decisions can still be impacted, for example, one study found that orthopedic surgeons had 22 times the odds of recommending total knee arthroplasty to a male patient as compared with a female patient.^51^ However, because our study controlled for patient-level demographic factors, and there does not appear to be literature specifically examining physician bias as it relates to insurance status, it is not clear whether physician bias based on insurance status may impact treatment decisions.

Another potential mechanism is unconscious or subtle pressures to reduce costs among this patient population, even if there do not appear to be policies or protocols that encourage this. There is a substantial cost to hospitals associated with treating uninsured trauma patients, with some studies suggesting that hospitals are unable to recoup costs associated with trauma care.^52-54^ Given that MRI scans are among the most expensive imaging procedures in the US,^55^ while plain X-ray radiography is a relatively low-cost technique, our finding of the strongest negative association between being uninsured and MRI use in both adjusted and IV analysis, and a positive association between being uninsured and X-ray use in IV analysis, are consistent with this mechanism.

A strength of our paper is that we used both standard adjustment and IV analysis, approaches which may be complimentary in that they have different potential biases. The fact that our findings were relatively consistent between these two different approaches strongly suggests an association between insurance status and imaging use, despite the vulnerabilities of each approach individually.

Our study used six years of data from the NTDB, a national data set that provided an adequate sample size for our analyses. However, our study was limited to patients who were treated at hospitals that report data to the NTDB, which may differ from other hospitals in that they may be generally higher resourced facilities with adequate infrastructure to comply with voluntary reporting procedures. If area-level SES interacts with patient insurance status, our findings may not be applicable to lower-resourced facilities than are represented by the NTDB.

Despite these limitations, our study provides evidence to support an association between insurance status and use of imaging, though the mechanisms for this remain unclear. Further quantitative and qualitative research could seek to elucidate potential reasons for the association found in our study. Another area of particular interest is whether disparities in hospital mortality related to insurance status can be in part addressed via improved diagnostics. Future studies should assess whether differences imaging use, or other diagnostic practices, mediate the frequently observed association between insurance and mortality.

## ACKNOWLEDGEMENTS

The National Trauma Data Bank is full and copyrighted property of The American College of Surgeons, Committee on Trauma. The American College of Surgeons Committee on Trauma is not responsible for any claims arising from works based on the original data, text, tables, or figures herein. No potential conflicts of interest (financial or otherwise) exist regarding this research for any of the authors.

## REFERENCES

1. Franks P, Clancy CM, Gold MR. Health insurance and mortality: Evidence from a national cohort. Jama. 1993;270(6):737–741.

2. Woolhandler S, Himmelstein DU. The relationship of health insurance and mortality: Is lack of insurance deadly? Annals of Internal Medicine. 2017;167(6):424–431.

3. Finkelstein A, Taubman S, Wright B, et al. The Oregon health insurance experiment: evidence from the first year. The Quarterly Journal of Economics. Aug 2012;127(3):1057–1106.

4. Van Der Wees PJ, Zaslavsky AM, Ayanian JZ. Improvements in health status after Massachusetts health care reform. The Milbank quarterly. Dec 2013;91(4):663–689.

5. Sommers BD, Gawande AA, Baicker K. Health Insurance Coverage and Health - What the Recent Evidence Tells Us. The New England journal of medicine. Aug 10 2017;377(6):586–593.

6. Statistics NCfH. Health, United States, 2016: With Chartbook on Long-term Trends in Health. 2017.

7. Taghavi S, Jayarajan SN, Duran JM, et al. Does payer status matter in predicting penetrating trauma outcomes? Surgery. Aug 2012;152(2):227–231.

8. Bell TM, Zarzaur BL. Insurance status is a predictor of failure to rescue in trauma patients at both safety net and non–safety net hospitals. Journal of Trauma and Acute Care Surgery. 2013;75(4):728–733.

9. Joseph B, Zangbar B, Khalil M, et al. Factors associated with failure-to-rescue in patients undergoing trauma laparotomy. Surgery. 2015/08/01/ 2015;158(2):393–398.

10. Haider AH, Chang DC, Efron DT, et al. Race and insurance status as risk factors for trauma mortality. Archives of Surgery. 2008;143(10):945–949.

11. Salim A, Ottochian M, DuBose J, et al. Does Insurance Status Matter at a Public, Level I Trauma Center? Journal of Trauma and Acute Care Surgery. 2010;68(1):211–216.

12. Lyon SM, Benson NM, Cooke CR, Iwashyna TJ, Ratcliffe SJ, Kahn JM. The Effect of Insurance Status on Mortality and Procedural Use in Critically Ill Patients. American Journal of Respiratory and Critical Care Medicine. 2011/10/01 2011;184(7):809–815.

13. Maybury RS, Bolorunduro OB, Villegas C, et al. Pedestrians struck by motor vehicles further worsen race- and insurance-based disparities in trauma outcomes: The case for inner-city pedestrian injury prevention programs. Surgery. 2010/08/01/ 2010;148(2):202–208.

14. Tepas JJ, Pracht EE, Orban BL, Flint LM. Insurance Status, Not Race, Is a Determinant of Outcomes from Vehicular Injury. Journal of the American College of Surgeons. 2011/04/01/ 2011;212(4):722–727.

15. Dozier KC, Miranda Jr MA, Kwan RO, Cureton EL, Sadjadi J, Victorino GP. Insurance Coverage Is Associated with Mortality after Gunshot Trauma. Journal of the American College of Surgeons. 3// 2010;210(3):280–285.

16. Rosen H, Saleh F, Lipsitz S, Rogers SO, Jr, Gawande AA. Downwardly mobile: The accidental cost of being uninsured. Archives of Surgery. 2009;144(11):1006–1011.

17. Zarzaur BL, Stair BR, Magnotti LJ, Croce MA, Fabian TC. Insurance type is a determinant of 2-year mortality after non-neurologic trauma. Journal of Surgical Research. 2010;160(2):196–201.

18. Englum BR, Villegas C, Bolorunduro O, et al. Racial, Ethnic, and Insurance Status Disparities in Use of Posthospitalization Care after Trauma. Journal of the American College of Surgeons. 2011/12/01/ 2011;213(6):699–708.

19. Nirula R, Nirula G, Gentilello LM. Inequity of Rehabilitation Services After Traumatic Injury. Journal of Trauma and Acute Care Surgery. 2009;66(1):255–259.

20. Zibulewsky J. The Emergency Medical Treatment and Active Labor Act (EMTALA): what it is and what it means for physicians. Proceedings (Baylor University. Medical Center). 2001;14(4):339–346.

21. Fowler RA, Noyahr L-A, Thornton JD, et al. An Official American Thoracic Society Systematic Review: The Association between Health Insurance Status and Access, Care Delivery, and Outcomes for Patients Who Are Critically Ill. American Journal of Respiratory and Critical Care Medicine. 2010/05/01 2010;181(9):1003–1011.

22. Bolorunduro OB, Haider AH, Oyetunji TA, et al. Disparities in trauma care: are fewer diagnostic tests conducted for uninsured patients with pelvic fracture? The American Journal of Surgery. 4// 2013;205(4):365–370.

23. Smith CB, Barrett TW, Berger CL, Zhou C, Thurman RJ, Wrenn KD. Prediction of blunt traumatic injury in high-acuity patients: bedside examination vs computed tomography. The American journal of emergency medicine. 2011;29(1):1–10.

24. Currie S, Saleem N, Straiton JA, Macmullen-Price J, Warren DJ, Craven IJ. Imaging assessment of traumatic brain injury. Postgraduate medical journal. Jan 2016;92(1083):41–50.

25. Looby S, Flanders A. Spine trauma. Radiologic clinics of North America. Jan 2011;49(1):129–163.

26. Parizel PM, van der Zijden T, Gaudino S, et al. Trauma of the spine and spinal cord: imaging strategies. European Spine Journal. Mar 2010;19(Suppl 1):8–17.

27. Vafaei A, Hatamabadi HR, Heidary K, Alimohammadi H, Tarbiyat M. Diagnostic Accuracy of Ultrasonography and Radiography in Initial Evaluation of Chest Trauma Patients. Emergency (Tehran, Iran). Winter 2016;4(1):29–33.

28. Raja A, Zane RD, Moreira ME, Grayzel J. Initial management of trauma in adults. 2012.

29. Bokhari F, Nagy K, Roberts R, et al. The ultrasound screen for penetrating truncal trauma. Am Surg. Apr 2004;70(4):316–321.

30. Rozycki GS, Ballard RB, Feliciano DV, Schmidt JA, Pennington SD. Surgeon-performed ultrasound for the assessment of truncal injuries: lessons learned from 1540 patients. Annals of surgery. Oct 1998;228(4):557–567.

31. Ho ML, Gutierrez FR. Chest radiography in thoracic polytrauma. AJR. American journal of roentgenology. Mar 2009;192(3):599–612.

32. Kumar Y, Hayashi D. Role of magnetic resonance imaging in acute spinal trauma: a pictorial review. BMC Musculoskeletal Disorders. July 22 2016;17(1):310.

33. Nathens A, Fantus R. National Trauma Data Bank 2010 annual report. Chicago, IL: American College of Surgeons Committee on Trauma; 2008.

34. Committee on Trauma ACoS. NTDB Annual Report 2015. 2015.

35. Freedman DA. On the so-called “Huber sandwich estimator” and “robust standard errors”. The American Statistician. 2006;60(4):299–302.

36. Hernán MA, Robins JM. Instruments for causal inference: an epidemiologist's dream? Epidemiology (Cambridge, Mass.). 2006;17(4):360–372.

37. Qayyum A, Yu JP, Kansagra AP, et al. Academic radiology in the new health care delivery environment. Academic radiology. Dec 2013;20(12):1511–1520.

38. Moser JW. The Deficit Reduction Act of 2005: policy, politics, and impact on radiologists. Journal of the American College of Radiology. 2006;3(10):744–750.

39. Thrall JH. Trends and developments shaping the future of diagnostic medical imaging: 2015 Annual Oration in Diagnostic Radiology. Radiology. 2016;279(3):660–666.

40. Azzalini L, Abbara S, Ghoshhajra BB. Ultra-low contrast computed tomographic angiography (CTA) with 20-mL total dose for transcatheter aortic valve implantation (TAVI) planning. Journal of computer assisted tomography. Jan-Feb 2014;38(1):105–109.

41. McGuire J, Wood BD. Prospective advancements in ultrasound imaging. Radiologic technology. Mar-Apr 2014;85(4):463–466.

42. Rassen JA, Schneeweiss S, Glynn RJ, Mittleman MA, Brookhart MA. Instrumental variable analysis for estimation of treatment effects with dichotomous outcomes. American journal of epidemiology. Feb 01 2009;169(3):273–284.

43. R: A language and environment for statistical computing. [computer program]. Vienna, Austria: R Foundation for Statistical Computing; 2016.

44. Kleiber C, Zeileis A. Applied econometrics with R. Springer Science & Business Media; 2008.

45. Jiang YS, D. ivpack: Instrumental Variable Estimation. R package version 1.2. https://CRAN.R-project.org/package=ivpack. 2014.

46. Goldfeld K. simstudy: Simulation of Study Data. R package version 0.1.3. https://CRAN.R-project.org/package=simstudy. 2017.

47. Bolorunduro OB, Haider AH, Oyetunji TA, et al. Disparities in trauma care: are fewer diagnostic tests conducted for uninsured patients with pelvic fracture? The American Journal of Surgery. 2013/04/01/ 2013;205(4):365–370.

48. Missios S, Bekelis K. The association of insurance status and race with the procedural volume of traumatic brain injury patients. Injury. 2016/01/01/ 2016;47(1):154–159.

49. Daly MC, Patel MS, Bhatia NN, Bederman SS. The Influence of Insurance Status on the Surgical Treatment of Acute Spinal Fractures. Spine. 2016;41(1):E37–E45.

50. Haider AH, Schneider EB, Sriram N, et al. Unconscious race and class bias: its association with decision making by trauma and acute care surgeons. Journal of Trauma and Acute Care Surgery. 2014;77(3):409–416.

51. Borkhoff CM, Hawker GA, Kreder HJ, Glazier RH, Mahomed NN, Wright JG. The effect of patients' sex on physicians' recommendations for total knee arthroplasty. Canadian Medical Association Journal. 2008;178(6):681–687.

52. Fath JJ, Ammon AA, Cohen MM. Urban trauma care is threatened by inadequate reimbursement. The American journal of surgery. 1999;177(5):371–374.

53. Henry MC, Thode HC, Shrestha C, Noack P. Inadequate hospital reimbursement for victims of motor vehicle crashes due to health reform legislation. Annals of emergency medicine. 2000;35(3):277–282.

54. Shen Y-C, Hsia RY, Kuzma K. Understanding the risk factors of trauma center closures: do financial pressure and community characteristics matter? Medical care. 2009;47(9):968.

55. Kliff S. How much does an MRI cost? In D.C., anywhere from $400 to $1,861. The Washington Post2013.

